# Dialogues in colour and behaviour - Integration of complex signalling traits and physiology

**DOI:** 10.1101/2024.08.16.608218

**Authors:** Subhasmita Patro, Thejaswini Saravanan, Ayush Parag, Maria Thaker

## Abstract

Animal communication can be complex, often involving multiple static and dynamic traits. The extent to which these traits are correlated can elucidate their function as either redundant or multiple messages. Using the agamid lizard, *Psammophilus dorsalis*, as a model system we examined patterns of trait expression and the role of steroid hormones in mediating these traits during social interactions. We staged male-male interactions in the lab and measured the repertoire of display behaviour and colours, which change dynamically in the visible and ultraviolet ranges in different body regions. Additionally, we measured testosterone and corticosterone levels before and after the social trials.

Our results show that within behaviour and colour trait categories, components were strongly correlated within individuals, suggesting either a shared physiological pathway or redundant information content. However, across trait categories, correlation patterns varied. The chromatic contrast of the (yellow) dorsal region of lizards was correlated with both body size and level of aggression, whereas the size of UV patches was correlated with body size only. We also found a negative association between baseline corticosterone levels, body size and dorsal yellow chromatic contrast, suggesting a mechanistic link between these traits. However, social interaction induced testosterone and corticosterone levels were uncorrelated with the expression of the dynamic behavioural and colour displays during the social interactions itself. Notably, the intensity of colour and behavioural displays of males were matched by their opponents. Overall, our results suggest that multiple signalling traits can ensure both redundancy as well as provide multiple messages to receivers, thus improving the robustness of information transfer, particularly during competitions which have high fitness consequences.

## Introduction

Social interactions in animals are often multi-faceted and complex. Such interactions can involve elaborate signalling, mediated by a combination of behavioural, morphological and physiological traits that interact in a complex feedback loop. While beneficial for communication, social interactions and their associated traits can also be costly. Maintaining multiple signals can be energetically costly to produce (McGraw et al. 2005, Stoddard and Salazar 2010) and, if conspicuous to predators or inter-specific competitors, can incur ecological costs to survival (Zambre et al. 2020; White 2022). Hence, given the potential costs, the use of multiple signals must be highly beneficial, and should be expected to enhance signal transmission, maintain signal honesty, or reduce receiver response time (Candolin 2003).

The evolution and maintenance of multi-component signals in animal communication has been explained using several broad frameworks of hypotheses. These are based on transmission efficacy, information content, and receiver psychology (Rowe 1999, Candolin 2003, Hebets and Papaj 2005). Efficacy-based hypotheses suggest that the signalling environment is variable and noisy and thus multiple signals evolved to ensure efficient transmission (Hebets and Papaj 2005). Content-based hypotheses assume that signals encode messages and thus provide information regarding signaller quality (Candolin 2003, Hebets and Papaj 2005). Within the content-based framework, the multiple messages hypothesis posits that animals use different signals to convey multiple pieces of information about the quality of the signaller (Hebets and Papaj 2005). Alternatively, the redundant signal hypothesis posits that multiple signals convey the same information, and these reinforce a single message (Hebets and Papaj 2005). However, not all signals provide information to receivers. Some signals are used to attract and orient receivers’ attention to a second signal. In the lizard *Amphibolurus muricatus*, tail flicking behaviour displayed at the start of an interaction serves to attract attention of the receiver (Peters and Ramos 2022). Signals can also act as amplifiers, where their presence increases the detectability of another signal (Hebets and Papaj 2005). Amplifiers and attention-grabbing signals are expected to be conspicuous, but less costly and uninformative (Castellano 2010). Lastly, receiver responses play an important role in signal evolution and maintenance, as having more than one signal is thought to increase detectability and reduce time and energy spent in assessing the signaller (Guilford and Dawkins 1991, Rowe 1999).

The complexity of social interactions is further augmented by multicomponent signals that can vary across time and space. Based on their temporal expression patterns, signals can be classified as static or dynamic. Signals such as body size, shape, or structure of weapons and ornaments are considered static or fixed, because they do not change in size or quality once developed. Many static signals reflect past conditions that animals have experienced and can act as honest indicators of individual quality. For example, body size, which is an outcome of conditions during development is an honest indicator of aggression during conspecific interactions in crayfish (Graham and Angilletta 2021). On the other hand, dynamic signals that can change during an interaction, reflect current conditions. Dynamic signals can be adjusted by the signaller depending on their internal state as well as receiver responses. Behaviour expressed during social interactions is a clear example of dynamic signalling (Partan and Marler 2005, Rosenthal 2007). Along with behavioural displays, several species belonging to taxa such as lizards, frogs and cephalopods can also dynamically change skin colours and patterns, to communicate (Hutton et al. 2015). Thus, static and dynamic signals can play different roles during social interactions.

Signals of similar types, such as colour or behaviour, displayed within a signalling context, can be causally linked through common mechanistic pathways. Similarity in structure or mode of production can lead to the encoding of similar information about the signaller (Shared pathway hypothesis – Index hypothesis, Weaver et al 2017). On the other hand, resource limitations may induce trade-offs between signals, resulting in expression of one signal at the expense of another (Resource trade-off hypothesis, Kodric-Brown and Brown 1984, Weaver et al. 2017). In Varied Tits (*Sittiparus varius*), carotenoid plumage colouration and tail length are negatively correlated, suggesting energetic trade-offs (Ma et al. 2023). Dynamic changes in skin colours on reptiles, amphibians and cephalopods are imparted by movements of pigment molecules within a chromatophore unit (reviewed in Ligon and McCartney 2016). Since they share the same biochemical pathways during signal expression, multiple colours might potentially encode redundant messages. On the other hand, different colour pigments such as carotenoids, melanin and structural UV colours are sourced and produced by different mechanisms in the body (Rawles 1948, Bagnara et al. 2007, Svensson and Wong 2010) and hence might reflect different types of information about the signaller’s foraging efficiency, immunocompetence, or dominance. Similarly, different behavioural displays require agile movement of body parts, involving muscles that need to endure high energy demands. The intensity of each type of behavioural display might therefore be limited by the motor capabilities of the relevant limb region. Thus, regardless of whether signals serve to attract attention or convey information, components of signals may be intrinsically linked because of their mechanism of expression.

What is not directly perceived by conspecifics during social interactions are the physiological mediators of signals. In many species, social interactions are known to elicit a change in steroid hormone levels that can affect signal expression. Numerous studies on rodents, lizards and birds have established the role of testosterone in facilitating heightened aggressive behaviour and expression of secondary sexual traits in males when faced with an intruder (Adkins and Schlesinger 1979, Wingfield et al. 1990). Not only does testosterone increase aggressive behaviour in males, but aggressive interactions themselves can increase plasma testosterone levels (Adkins-Regan 2005). The association between testosterone and aggression, however, is greatly dependent on the social and ecological context, season, age, experience and taxa (Wingfield et al. 1987) and hence is subject to great variability. Steroid hormones have additional roles in the communication context. In some species, testosterone has been shown to increase the bioavailability of carotenoids (Blas et al. 2006, Khalil et al. 2023), due to its role in immunosuppression (Immunocompetence handicap hypothesis; Folstad and Karter 1992). Additionally, since social interactions can be energy intensive, glucocorticoid hormones are also expected to influence individual responses. Rise in glucocorticoids during social challenges redirects energy towards immediate needs (Wingfield and Romero 2001, Sapolsky et al. 2000). Glucocorticoids are also known to affect melanin-based colouration (Roulin 2008) by inhibiting melanogenesis (Slominski 2004), and when chronically elevated, can redirect carotenoids from ornaments to immune functions (Loiseau 2008).

Since the expression of signals can be affected by any or all the above factors, unravelling the relationship between multiple signals and their underlying physiology remains a challenge. We determined the patterns of signal correlation using the tropical agamid lizard, Indian rock agama (*Psammophilus dorsalis*). During social interactions, males of this species display dynamic colour change (Batabyal and Thaker 2017), that involve areas of their skin that is pigmented with carotenoid- and pterin-based red, orange, and yellow patches (Amdekar and Thaker 2022), melanin-infused brown and black, as well as ultraviolet patches. Changes in the spectrum of colour in males can occur within seconds of exposure to a conspecific male or female (Batabyal and Thaker 2017). When males interact with other males, rapid colour change is accompanied by display behaviours that are directed towards the competitor (Deodhar and Isvaran 2018, Batabyal and Thaker 2017, 2019). Previous studies have shown that social interactions in male lizards can elicit an increase in testosterone and corticosterone levels, which stay elevated for 10-30 min after the interaction (Batabyal and Thaker 2019). Thus, the Indian rock agama uses a combination of static and dynamic signals comprising of colour, behaviour and morphological traits to communicate during social interactions. How these multiple signalling traits interact and whether they influence receiver responses are still unclear. In this study, we quantified patterns of correlation between male body size, display colours, behaviours, and hormone levels. Patterns of correlation between trait components will indicate the degree of signal redundancy. We also examined the correlation between these signal components and both baseline and interaction-induced testosterone and corticosterone levels to identify potential mechanisms involved in the expression of signals. Finally, we assess whether receiver responses are influenced by one or more of these signal components. Overall, our study enables us to evaluate the potential hypotheses for multi-component signals in a system that has evolved both static and dynamic traits for animal communication.

## Methods

### Field capture and housing

Adult male lizards from the wild (N=55) were caught by lassoing during the months of June to September 2021, from 4 study sites on the outskirts of Bangalore, that were 5-10km apart. After capture, lizards were placed in separate cotton bags and transported in a cooler lined with icepacks to the lab, where they were housed individually in glass terraria (60 × 30 × 25 cm). Terrariums were lined with paper towels and provisioned with rocks and a shelter for basking and refuge. Terrariums were maintained in a dedicated lizard housing room, which was equipped with automated 12-hour light/dark cycle and maintained at ambient temperature conditions. Individual terrariums were also set up with incandescent basking lamps (60W) that were turned on for 5 hours during the day. Lizards were provided with mealworms and grasshoppers daily for food, along with *ad libitum* water. All lizards were allowed to acclimate for three days to laboratory conditions before the start of trials and were maintained in the lab for 9-10 days for the experiments, following which they were released at their site of capture.

### Blood Sampling and Hormone Analyses

We collected two blood samples from every individual. To determine baseline circulating hormone levels, ∼50µL of blood was drawn from the retro-orbital sinus of each lizard using a heparinized microcapillary tube (Batabyal and Thaker 2019) after three days of acclimation in the laboratory. Lizards were allowed two days to recover from the first blood sample, before behavioural trials began. To measure social interaction-induced hormone levels, a second sample of ∼50µL of blood was drawn, immediately at the end of the staged behavioural trials (see below). Immediately after collection, blood samples were centrifuged and the plasma stored at −20°C, until hormone analysis. Enzyme-Immuno Assay kits (Arbor Assay DetectX Corticosterone K014-H5; Testosterone K032-H5), optimized for the species (Batabyal and Thaker 2019), were used to measure concentrations of circulating testosterone and corticosterone levels in the plasma. We analysed plasma at a dilution ratio of 1:100 or 1:80 for corticosterone and 1:140 or 1:160 for testosterone. For both hormones, samples were run in triplicates and a total of 13 assays (plates) were run, with a duplicate lab standard in each plate. For corticosterone, mean intra-assay coefficient of variation was 8.08 (ranging from 0.07-17.6) and inter-assay coefficient of variation was 15.78. For testosterone, mean intra-assay coefficient of variation was 8.38 (ranging from 1.05 to 18.1) and inter-assay coefficient of variation was 15.07.

### Social interaction Trials

We measured the snout-vent length (SVL) of all lizards, so that they could be size-matched for the social interaction trials. Individuals (N=46) in a pair were selected from different field sites such that they were expected to be unfamiliar with each other. Trials were staged in a large glass terrarium (100×40×30cm) in the lab. Lizards were introduced into the testing terrarium on either side of an opaque divider and allowed 60 minutes to habituate. Once the divider separating them was removed, the pair was allowed to interact for 30 minutes. The experimental area was lit using four fluorescent tube lights, one UV tube light (39 watts, Reptisun 10.0 UVB T5, Zoo Med Laboratories Inc) and a full spectrum bulb (VivaLite: B22).

Control trials (N=9) were staged with a single lizard in the experiment tank with all other conditions remaining the same. All trials along with blood sampling for hormone analysis took place between 9am to 12:30pm, which coincides with the peak activity hours of *P. dorsalis*.

Lizards in social interaction and in control trials were digitally recorded on video as well as photographs, using two cameras simultaneously. To enable the quantification of behaviour, digital videos were recorded using a Cannon 700D camera with an 18-135 mm lens. To obtain images of body colouration in the UV and visible range, we took digital photographs of each lizard at 4-minute intervals using a modified multi-spectral Nikon D5500 camera. This camera was fitted with a Nikkor EL 80mm lens, and either a 250-390 nm filter for the UV images (Fotodiox) or 400-750 nm filter (Fotodiox) for visible only images. Photograph of a colour checker (X-rite model: MSCCPP) under the same lighting conditions was used to standardize the colours in the visible range. Teflon tape placed in the tanks was used as reference for UV photographs.

### Social behaviour

From the digital recordings of the 30 min social interactions, we quantified behaviours displayed (see Table 1) by both the males of an interacting pair (similar to Batabyal and Thaker 2019).

**Table 1.**
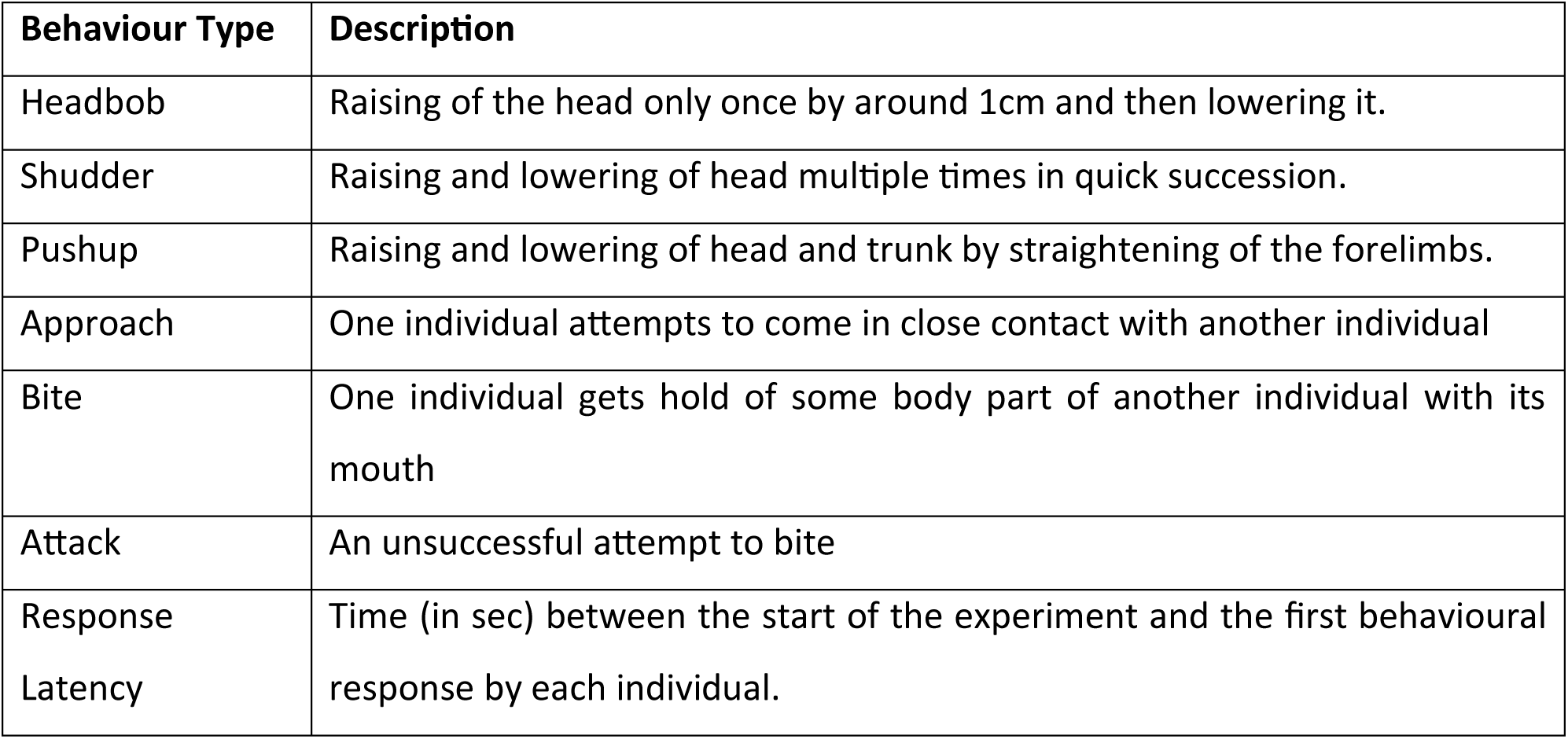
Behaviours displayed by males during the staged male-male interactions. All behaviours were quantified from videos.

All behaviours except response latency were measured as events (i.e. count). For statistical analyses, we combined the bite and attack responses (hereafter known as BiteAttacks), since the occurrence of these two behaviours was rare in all interactions.

### Skin Colour Measurement

We had 6-10 digital images in the visible-only range of each lizard during the trials. Skin colour was quantified from these images as chromatic contrasts measured in terms of Just Noticeable Differences (JND), using the MicaToolBox plugin in ImageJ (van den Berg et al. 2019). All images were first linearized using five grey scale reflectance squares on a standard colour checker (X-rite model: MSCCPP) and a multispectral image was created. In the linearized multispectral image, we created a 2×2mm square region of interest (ROI) each on the dorsal and the lateral body region of the lizard (Supplementary Figure S1). Using the inbuilt chameleon eye model (with cone ratios set as lw:mw:sw = 0.06:0.08:0.15) and daylight (D65) lighting in the MicaToolBox, we extracted the cone catch values for these ROIs. The cone catch values were then converted to the receptor noise limited (RNL) colour space, and chromatic contrasts were calculated against a standard 40% grey background. The maximum chromatic contrast on both dorsal and lateral body region, displayed by each individual during the 30-minute interaction period was considered for further analysis. We also measured the chromatic contrast between the dorsal yellow and the lateral orange colour (henceforth: internal chromatic contrast).

### UV Patch Area Measurement

UV patch area or brightness was not measured from images taken during the trials, because lizards were free to move and therefore their bodies were in different angles with respect to the camera, leading to an under-estimation of patch size. Instead, two days after the behavioural trials, we induced a stress response in males by restraining each lizard for 10 min. Restraint stress reveals the maximum size of the UV patches, which we then photographed (Supplementary Figure S2) using the Nikon D5500 multi-spectral camera with a 250-390 nm filter on the lens.

From these images, we measured the area of the UV patches on the lizard body using the image segmentation technique in ImageJ v1.53K. We employed a slightly modified protocol used for data acquisition from images by Badiane and Font 2021 and Garcia et al. 2013. The UV patches were separated from the rest of the lizard body using the image segmentation tool, “threshold”, with manual thresholding under the “Shanbhag” mode (Figure S2). Because individuals differ in the intensity of UV patch brightness and the size and shape of the UV patches on their body, manual thresholding was used to avoid over or under estimation of the UV-covered area. The images were then cleaned using the paintbrush tool to remove any false patches (such as shiny scales or shed skin) that may appear as UV regions. Following thresholding and cleaning, an ROI was created, and the total area of interest was measured by using the “Analyse Particles” tool. Every image had a graph sheet in it that was used as a reference for scale.

### Statistical Analyses

We first tested the normality of every measured trait using Shapiro Wilk’s test, and accordingly used either parametric or non-parametric tests for further comparisons. Blood samples were obtained between 71 to 431 seconds from the end of social interactions. However, there was no significant effect of blood sampling time and any of the hormone levels (p<0.05 for all comparisons). Three individuals in the social interaction trials that had extremely high testosterone levels (>2000ng/ml) were excluded from all further hormone analyses.

To determine differences between males in control trials and those in social interactions, we compared hormone levels with a Wilcoxon rank sum test, and chromatic contrasts of colour patches with Welch’s two sample t-test for the dorsal yellow region and Wilcoxon rank sum test for the lateral orange region. For the social trials, we also tested the difference between baseline and interaction-induced hormonal responses with a Wilcoxon paired sample test.

When determining the correlations between traits during the social trials, we first reduced the behavioural responses using a Principal Component Analysis into PC1 (62% of the data explained) and PC2 (22%) (See Supplementary Figure S3b). All pair-wise correlation tests between traits were performed using either Pearson’s or Spearman’s correlation test depending on whether the data were normally distributed (SVL, mass and dorsal yellow chromatic contrast) or non-normally distributed (testosterone level, corticosterone level, lateral orange chromatic contrast, and PC1 and PC2 of behavioural responses).

Since the social interactions were paired and we collected data from both the individuals involved, every individual in this study is a signaller as well as a receiver (“opponent”). We performed pair-wise correlation tests (either Pearson’s or Spearman’s) between individual responses and opponent responses for all colour and behaviour traits.

## Results

### Hormonal Responses

Following the 30-minute social interaction, males showed a significant increase in corticosterone level compared to baseline (Wilcoxon paired sample test: V = 293, p-value=0.01). But there was no significant difference between baseline and interaction-induced testosterone levels (V = 356, p-value = 0.24). Testosterone (Wilcoxon rank sum test: W = 250, p-value = 0.14) and corticosterone (W = 215, p-value = 0.86) levels of individuals in control trials (no social interaction) and social trials did not vary significantly at the end of the trials (30 minutes).

### Behavioural Responses

During staged male-male encounters, 40 out of 46 animals displayed at least one behaviour. Headbob and shudder were the most common behaviours displayed (Figure S2a). In every trial that elicited a behavioural response, the encounter started with a headbob or shudder and then the pair of males aligned themselves parallel to each other. The encounter then either escalated into an attack and bite (41%) or more vigorous head bobs (59%). Two individuals that displayed very high numbers of shudder displays were excluded from the statistical analyses. As expected, animals in the control trials did not show any change in behaviour for the duration of the experiment.

When reducing these behavioural responses with a PCA, all behaviours loaded positively on PC1, with loadings ranging from 0.36 to 0.49. Headbobs (−0.69) and shudders (−0.46) loaded negatively on PC2, whereas all other behaviours loaded positively (0.29 to 0.35) on PC2. Bartlett’s test of sphericity on the PCA yielded chisq=117.83, p <<<0.001, suggesting that PC1 and PC2 represent most of the variation in the data.

### Physiological colour change

All animals displayed some dynamic colour change during the interaction. Overall, animals in the social interaction trials showed higher chromatic contrasts in both dorsal (Welch two sample t-test: t = 2.46, p-value=0.03,) and lateral body regions, (Wilcoxon rank sum test: W = 306, p-value = 0.02) compared to control animals (See Supplementary Figure S1). Additionally, chromatic contrast of the dorsal region was higher than the lateral body region (Wilcoxon rank sum test: W = 472, p<0.001,) during social interaction. Moreover, internal chromatic contrast between the dorsal and lateral regions in the social males was lower than the individual contrasts of these regions measured separately (See Supplementary Figure S4).

### Correlation <u>within</u> Hormone levels, Behaviour and Colour traits

Testosterone levels induced during the social interaction were positively correlated with baseline levels (r = 0.78, p < 0.01). Baseline and social interaction-induced levels of corticosterone were also positively correlated (r = 0.72, p < 0.01). However, baseline testosterone was not correlated with baseline corticosterone levels (r = 0.07, p = 0.65, Figure S6a) and neither were interaction-induced testosterone and corticosterone levels (r = −0.09, p = 0.57, Figure S6b).

Most behavioural traits displayed during social interaction were positively correlated with each other (r > 0.32, p <= 0.037) and negatively correlated with response latency (r < −0.21, p <= 0.05, see Supplementary Table S1).

Similarly, the maximum chromatic contrasts of dorsal yellow and lateral orange regions were also positively correlated within individuals (r = 0.38, p = 0.01, Table S2). However, the area of UV patches was not correlated with the other colour measures (r < 0.23, p > 0.1, Table S2).

### Correlation <u>between</u> Behaviour, Colour and Hormone levels

Behaviour (PC1) displayed during social interaction was positively correlated with the maximum dorsal yellow chromatic contrast shown during the interaction (r = 0.44, p-value = 0.003; Figure 3), but not correlated with any of the hormone measures (Figure 2).

**Figure 1.**
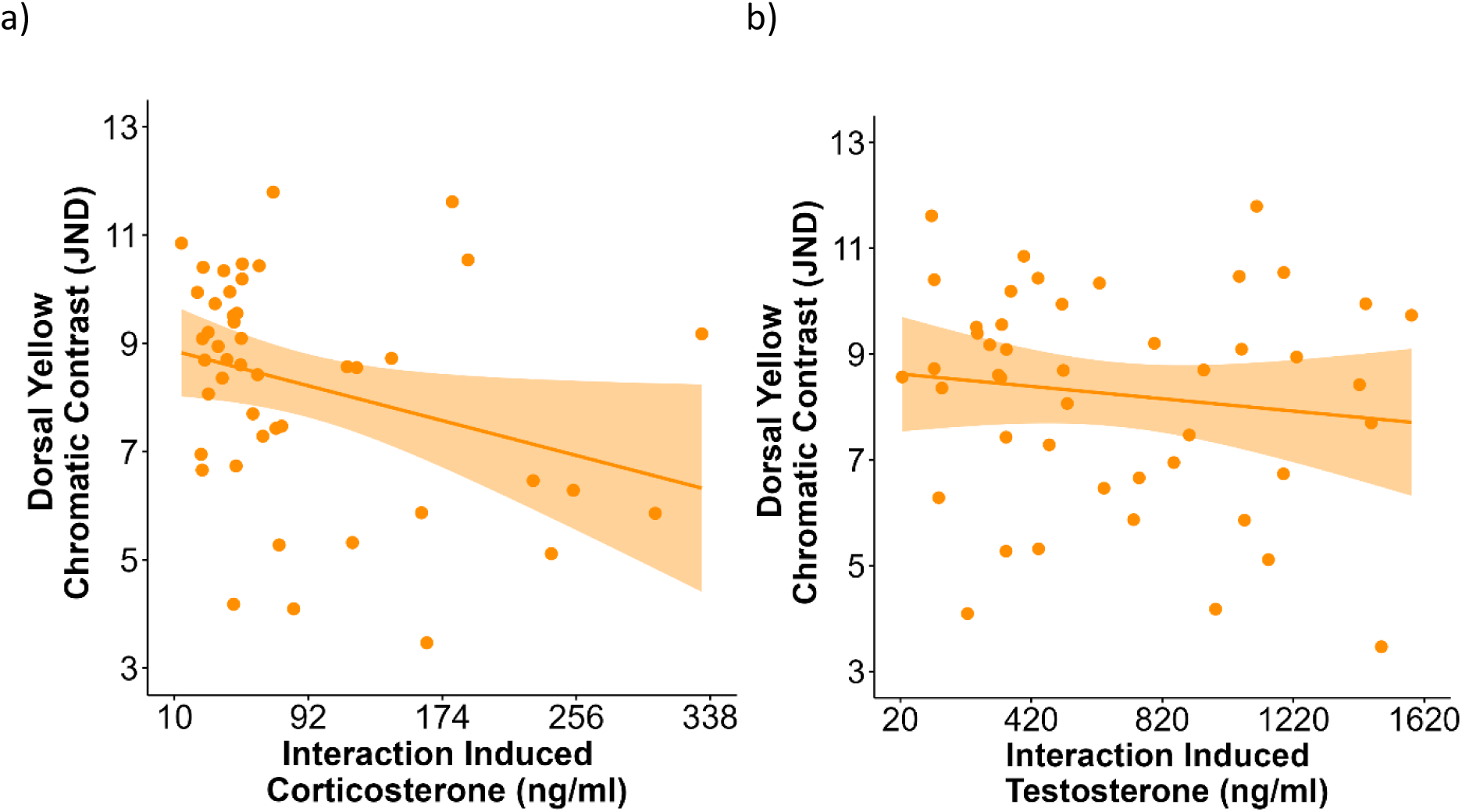
(a) Interaction-induced levels of corticosterone was negatively correlated with the chromatic contrast in the dorsal body region (r = −0.33, p = 0.03), but (b) there was no significant correlation between interaction-induced testosterone level and dorsal colour (r = −0.1, p = 0.5).

**Figure 2.**
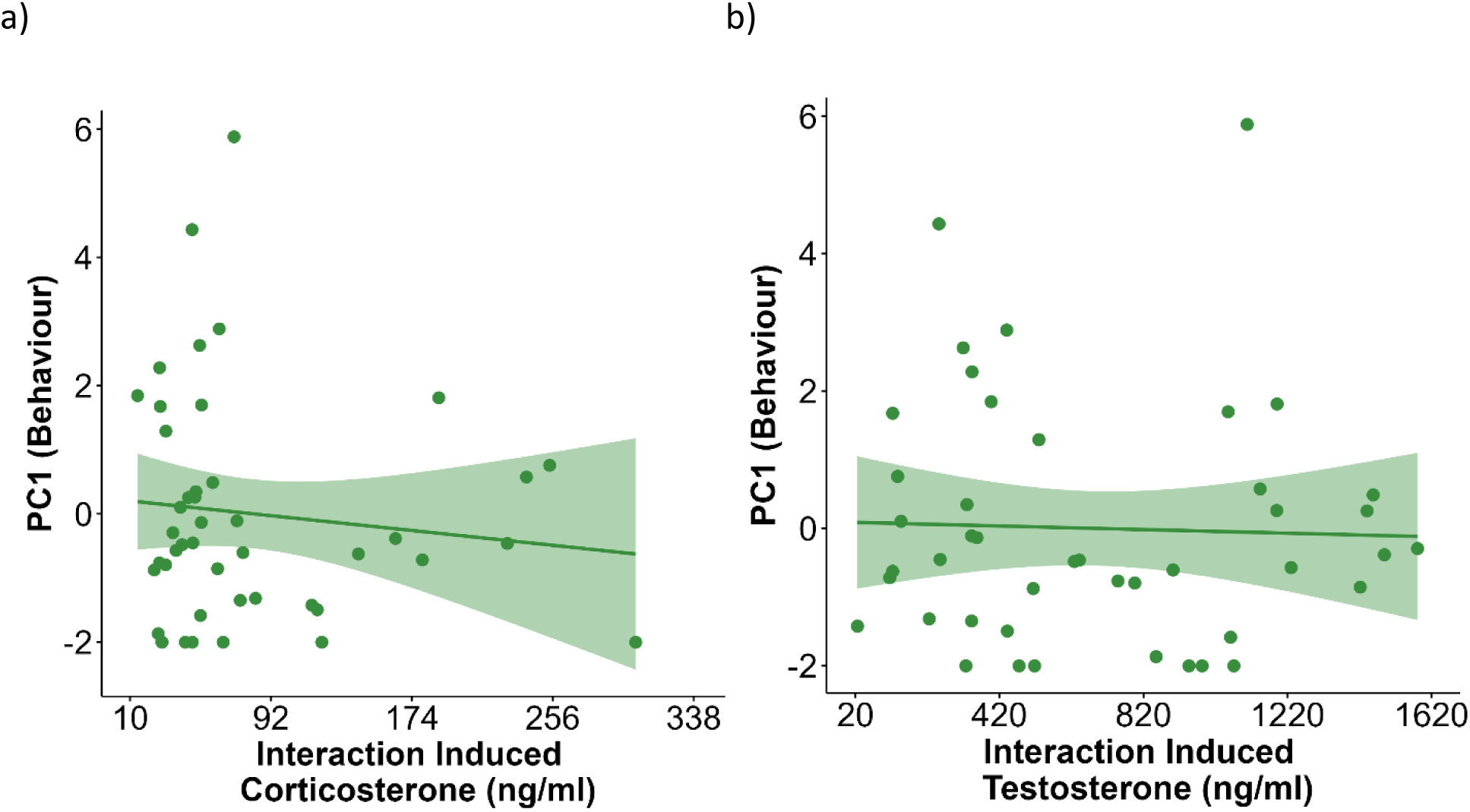
Interaction induced levels of (a) corticosterone (r = −0.076, p = 0.62) and (b) testosterone (r = 0.03, p = 0.85) were not correlated with behaviour displayed during social interaction.

**Figure 3.**
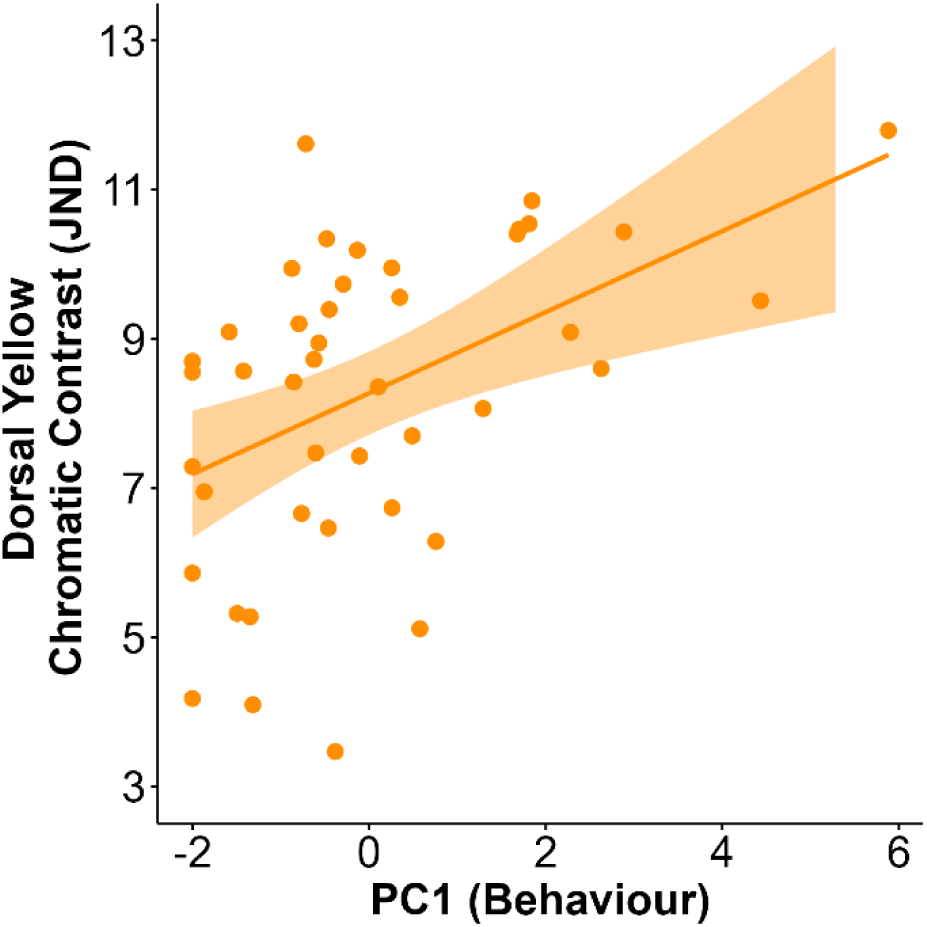
Behaviours displayed during social interaction was positively correlated with dorsal yellow colour expressed by *P. dorsalis* during male-male social interactions (r = 0.44, p = 0.0031)

Both baseline and interaction-induced corticosterone levels were negatively correlated with the maximum dorsal yellow chromatic contrast in males (baseline: r = −0.41, p-value = 0.005; Figure S5a; interaction-induced: r = −0.33, p-value = 0.03, Figure 1a). Corticosterone levels were not correlated with any other colour measures. None of the colour measures were correlated with baseline (Figure S5b) or interaction-induced testosterone levels (Figure 1b).

Body size (snout-vent length, SVL) was negatively correlated with baseline corticosterone (r = −0.38, p-value = 0.01; Figure 4a) but was uncorrelated with all other hormone measures (Figure 4b for baseline testosterone, Figure S7 for rest). SVL was also positively correlated with the maximum dorsal yellow chromatic contrast (r = 0.42, p-value = 0.004; Figure 4c) and internal chromatic contrast (r = 0.29, p-value = 0.05) displayed during the interactions as well at the total UV patch area (r = 0.45, p-value = 0.003; Figure 4d).

**Figure 4.**
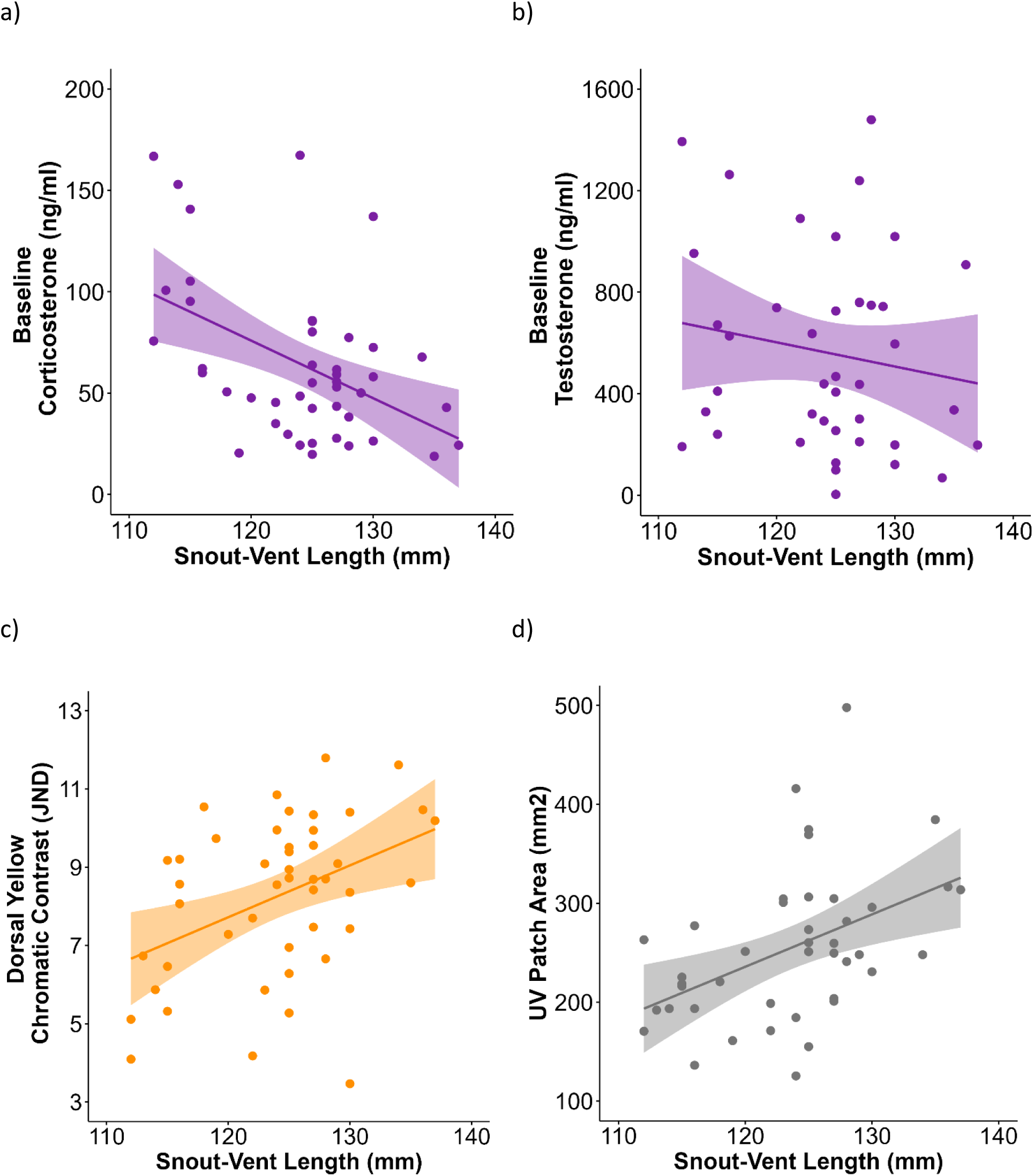
Body size (snout-vent length) was negatively correlated with a) baseline corticosterone level (r = −0.38, p = 0.01) but uncorrelated with b) baseline testosterone level (r = −0.11, p = 0.48). SVL was positively correlated with c) maximum chromatic contrast of dorsal yellow colour (r = 0.42, p = 0.004) and d) total area of UV patches in social males (r = 0.45, p = 0.003).

### Opponent Response

Both the display behaviour and dorsal chromatic contrast of an individual was positively influenced by the opponent’s dorsal colour (behaviour: r = 0.46, p-value = 0.002; colour: r = 0.52, p-value < 0. 001; Figure 5a) and behaviour (behaviour: r = 0.56, p-value < 0.001; colour: r = 0.51, p-value < 0.001; Figure 5b).

**Figure 5.**
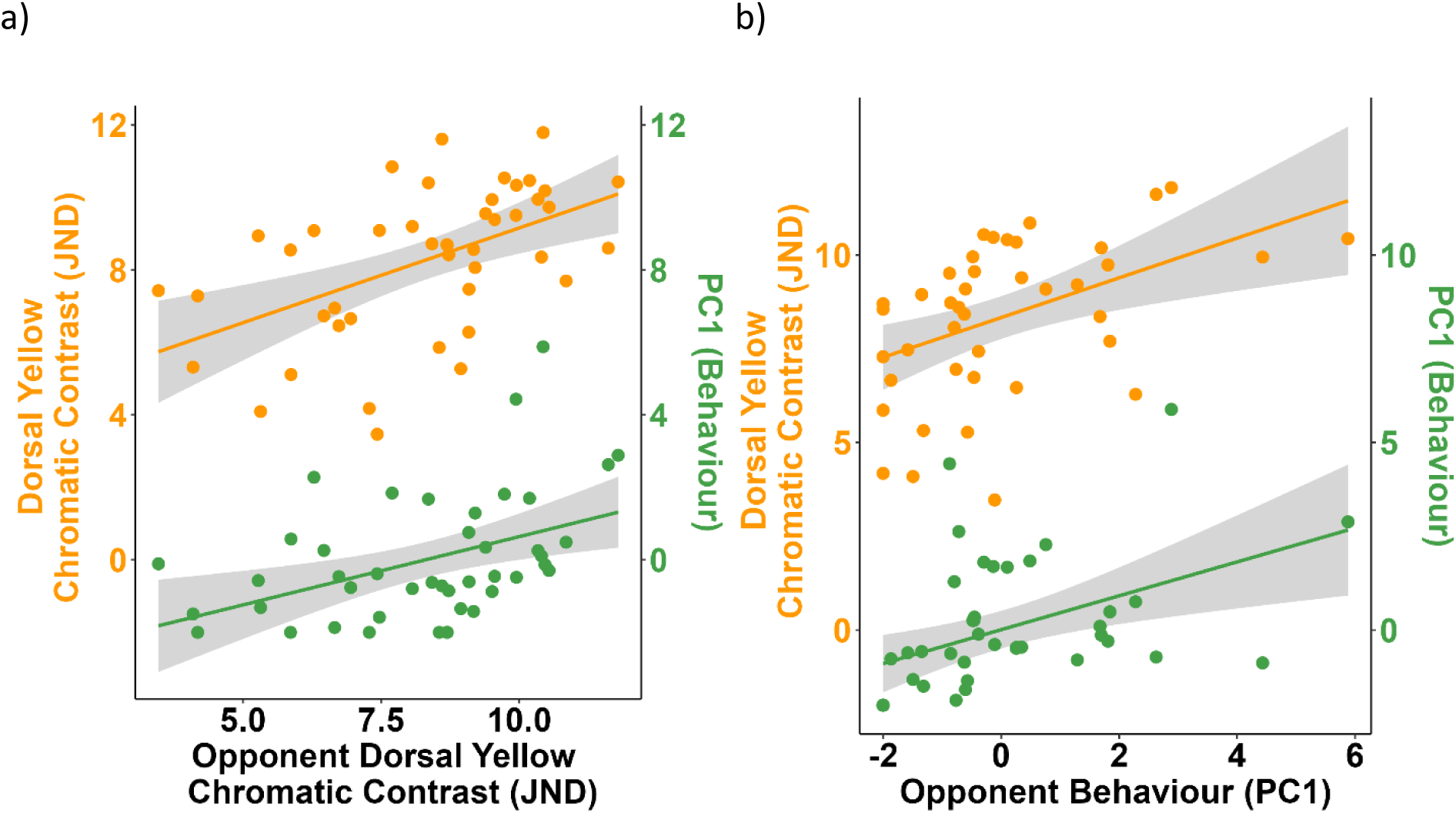
Behaviours (green) and dorsal yellow colours (orange) displayed during a social interaction are positively influenced by the opponent’s a) Dorsal yellow chromatic contrast and b) Behaviours (PC1).

## Discussion

A complex signalling system is generally composed of multiple components that either convey redundant information, encode multiple messages, or are attention-grabbing. Components of the signalling system can therefore be coupled (correlated) or decoupled (uncorrelated) depending on their function or mechanism of expression. In *P. dorsalis*, we find that social interactions between males involve the expression of multiple behavioural and colour displays, and an elevation of corticosterone but not testosterone. Within the categories of colours, behaviours and hormones, the multiple traits expressed are highly correlated within individuals. However, there were only a few correlations across trait categories, indicating that the expression of colours, behaviours, and hormone levels during social interactions are fairly independent of each other.

Signals that contain the same information (i.e. are redundant) or that share the same pathway of expression are expected to be strongly correlated. In *P. dorsalis*, we found positive correlations within the behavioural displays and within some of the colours expressed. Animals that displayed more headbobs, shudders and pushups were also more likely to approach, attack and bite. Variation in overall behavioural activity may indicate inherent differences among males in the level of aggression that they express during male-male interactions, consistent with observations in other lizard species (Baird et al 2013, Payne et al 2021). Shared physiological pathways could explain the correlation of behaviours within individuals, as the expression and intensity of behaviours is largely dependent on the musculo-skeletal endurance of the limbs and neck as well as motor capabilities (Higham et al. 2011, Albuquerque 2023). Among the colours expressed, both the dorsal yellow and lateral orange colours in *P. dorsalis* are composed of multiple pterins and carotenoid pigments (Amdekar and Thaker 2022), and thus a shared biochemical pathway may explain the strong correlation in the chromaticity of these colours during social interactions. Notably, males also express UV reflective patches during aggressive and stressful situations, similar to other species (Whiting et al 2006). Yet, we find the patch size of these dynamically expressed UV patches to be uncorrelated with the intensity of chromatic colours, indicating that pigment and UV colours do not have a shared pathway and are potentially non-redundant.

Since conspicuous colours and behaviours are easily discernible by receivers, but are potentially costly to produce and maintain, these traits could signal individual quality. Numerous studies on birds, fish and reptiles have reported the usage of carotenoid, UV and melanin colours to signal aggression and body size during social contests (Cook et. al. 2013, Kemp and Grether 2014). In *P. dorsalis*, we find that individuals that have higher dorsal yellow chromatic contrast are also more aggressive and have larger body size. However, contrary to our expectations, the chromatic contrast of the lateral orange colour was uncorrelated with behaviour or body size. Previous studies with this species have shown that the chromatic contrasts of the dorsal yellow and lateral orange colours are not associated with sprint speed and bite force in males (Amdekar and Thaker 2022). Hence, it is possible that the lateral orange colour might act as an amplifier or attention-directing signal, whereas the yellow dorsal colour region is where information about behaviour and morphology are encoded. As defined in Hebets and Papaj 2005, an attention directing signal quickly captures the receiver’s attention and alerts them to a subsequent information-containing signal. Thus, the attention directing signal is expected to be more conspicuous than the information-containing signal. Similarly, an amplifier signal is expected to lower the detection threshold of the second signal, making the combined signal more conspicuous, while each signal alone is difficult to detect. However, in *P. dorsalis*, the chromatic contrast of the lateral orange is lower than the chromatic contrast of the dorsal yellow, indicating that the yellow colour would be more readily visible to a receiver compared to the orange. Furthermore, if the orange colour acted as an amplifier, then the internal chromatic contrast between the yellow and orange should be higher than the individual chromatic contrasts (Hebets and Papaj 2005). However, we do not see this pattern (Figure S1), and thus the amplifier function of the lateral orange colour is unlikely. It is possible that the lateral orange colour might just be an involuntary physiological response with no signalling role. Lastly, UV colours have been used as badges of status and to signal aggressiveness in many lizard species (Whiting et al. 2006, Martin et al. 2016). In *P. dorsalis*, UV reflective patch sizes on the dorsal region of males correlated strongly with body size but not with other measures of behaviour or colour. Given that UV patches are dynamically revealed during stress and aggression, males can reliably assess each other’s body sizes using UV patch sizes as honest indicators.

In many vertebrates, testosterone is associated with modulation of the expression of signals, such as colours and behavioural displays, during social interactions (Adkins-Regans 2005). For androgens to directly modulate social behaviours or dynamic colour expression, we would expect individuals to increase testosterone levels during social encounters (i.e. Challenge hypothesis, Wingfield et al. 1990). However, evidence from a growing body of studies on tropical species does not always comply with this pattern (Roberts et al. 2003, Lynn 2008). Social interactions in *P. dorsalis* did not elicit an increased testosterone response in our study, which is similar to findings in several other species of birds, fish and lizards (Cramer 2012, Ros et al. 2014, Ligon and McGraw 2018). Additionally, testosterone levels did not correlate with a male’s display colour or behaviour. Since these experiments were conducted during the breeding season and animals were likely already involved in breeding-related social interactions before capture, it is possible that testosterone levels were sufficiently elevated to facilitate an aggressive response, a pattern widely observed in many tropical species (Wingfield et al. 1987, Wingfield et al 1990, Moore et al. 2019).

Following social interactions in *P. dorsalis,* we detected a rise in plasma corticosterone from baseline levels. Both baseline and interaction-induced corticosterone levels were negatively correlated with the chromatic contrast of a male’s dorsal yellow colour during social interactions. Corticosterone levels were uncorrelated with other measures of colour or behaviour. Although evidence linking glucocorticoids to ornamentation is mixed (Leary and Baugh 2020, Moore et al. 2016), our findings support multiple studies in birds, reptiles and amphibians that have found negative associations between baseline glucocorticoid levels and carotenoid-based colours (Loiseau et al. 2008, San-Jose and Fitze 2013, Gardner et al. 2020). Both baseline corticosterone levels and dorsal yellow colour were also correlated with the body size (SVL) of the lizards, such that larger bodied males had lower levels of plasma corticosterone and higher contrast of dorsal yellow. This pattern suggests a physiological link between baseline corticosterone, body colour (dorsal) and body size. Since dorsal colours are also indicators of aggressive behaviour in *P. dorsalis*, corticosterone might play a mechanistic role in reinforcing signal redundancy and honesty in this species, similar to findings in other studies (Tibbetts 2014, Vitousek et al. 2014). Although we expected corticosterone levels to influence aggressive behaviour directly (Summers et al. 2005), we find no support for this in our data. Since males who did not engage in a social interaction (control individuals), also showed a rise in corticosterone from baseline levels over the same duration, it is possible that 30 minutes in a novel environment elicits a stress response regardless of the additional social encounter. Alternatively, corticosterone could also have a permissive role, where its presence is necessary only for expression but not modulation of aggressive behaviour (Sapolsky 2000).

Receiver response is widely accepted as one of the major factors that drive signal evolution and maintenance (Tibbetts 2014). However, the ability to modulate signal quality leaves ample room for signal manipulation and dishonest signalling (Dawkins and Guildford 1991, Wilson and Angilletta 2014). For example, male mourning cuttlefish (*Sepia plangon*) tactically display deceptive signalling patterns to prevent rival males from gaining courtship (Brown et al 2012). Nevertheless, aggressive interactions with receivers are known to impose a cost that ensures signal honesty over evolutionary time (Tibbetts 2014). We found that during male-male interactions with size matched opponents, behavioural displays and dorsal colours expressed by *P. dorsalis* males are strongly affected by the signaller’s behaviour and dorsal colour. This means that males with high chromaticity of dorsal colour and high behavioural aggression elicited similarly high responses in both these traits in their opponents. Receiver response might reinforce signal honesty in this system, since conspicuous dorsal colours and aggression elicit more aggression and attacks from the opponents, making it difficult for weak individuals to cheat. Note that males also express UV patches that are correlated with body size, and thus UV patch size may also be an important signal of honesty if opponents were mismatched in size.

Overall, the complex signalling system of *P. dorsalis* involves the expression of many traits. Our results show strong correlation within similar trait types (colour, behaviour, hormones) indicating a shared physiological pathway or redundancy in information content. Additionally, we find evidence for multiple messaging as well as redundancy across some trait components. While the chromaticity of the dorsal yellow colour and UV patch sizes provide information on male body size, the level of behavioural aggression is indicated by the dorsal yellow colour. The maintenance of these potential signals is further reinforced by the receiver’s response to them. However, the role of lateral orange colour in male-male aggression is still not known. Our findings also point towards a link between body size, chromaticity of dorsal colour and baseline corticosterone level, providing further support for the maintenance of honesty of a dynamic colour signal. Contrary to our expectations, we found no link between testosterone level and aggressive behaviour or colour. Similar lack of association between corticosterone and behaviour was also unexpected, indicating possible permissive roles for both the hormones in the context of male-male interactions. Overall, the presence of complexity in a signalling system underscores the need for both redundancy and non-redundancy, especially during male-male interactions that have fitness consequences.

