## Supplement File 1 for "Dialogues in colour and behaviour - Integration of complex signalling traits and physiology"

### Supplementary Material

#### Tables and Figures

**Table S1:** Behaviours displayed by males of *P. dorsalis* during staged male-male interactions. Shown are correlation coefficients (r) and p-values from spearman's correlation test. * represents p <= 0.05.

| | **HeadBob** | **Shudder** | **Pushup** | **Approach** | **BiteAttacks** |
|---|---|---|---|---|---|
| **Response Latency** | **r=-0.52, p=0.0003*** | **r=-0.50, p<0.001*** | r=-0.21, p=0.174 | **r=-0.36, p=0.015*** | **r=-0.29, p=0.055*** |
| **HeadBob** | | **r=0.77, p<<0.001*** | **r=0.32, p=0.037*** | **r=0.40, p=0.007*** | **r=0.32, p=0.036*** |
| **Shudder** | | | **r=0.56, p<<0.001*** | **r=0.55, p<0.001*** | **r=0.4, p=0.007*** |
| **Pushup** | | | | **r=0.72, p<<0.001*** | **r=0.66, p<<0.001*** |
| **Approach** | | | | | **r=0.79, p<<0.001*** |

**Table S2:** Maximum chromatic contrast of the colours displayed by males of *P. dorsalis* during staged male-male interactions. Shown are correlation coefficients (r) and p-values from spearman's correlation test. * represents p <= 0.05.

| | **Lateral Orange Chromatic Contrast** | **Internal Contrast** | **Total UV Area** |
|---|---|---|---|
| **Dorsal Yellow Chromatic Contrast** | **r=0.38, p=0.01*** | **r=0.77, p<<0.001*** | r=0.18, p=0.25 |
| **Lateral Orange Chromatic Contrast** | | **r=0.39, p=0.009*** | r=-0.08, p=0.61 |
| **Internal Contrast** | | | r=0.23, p=0.14 |

**Figures**

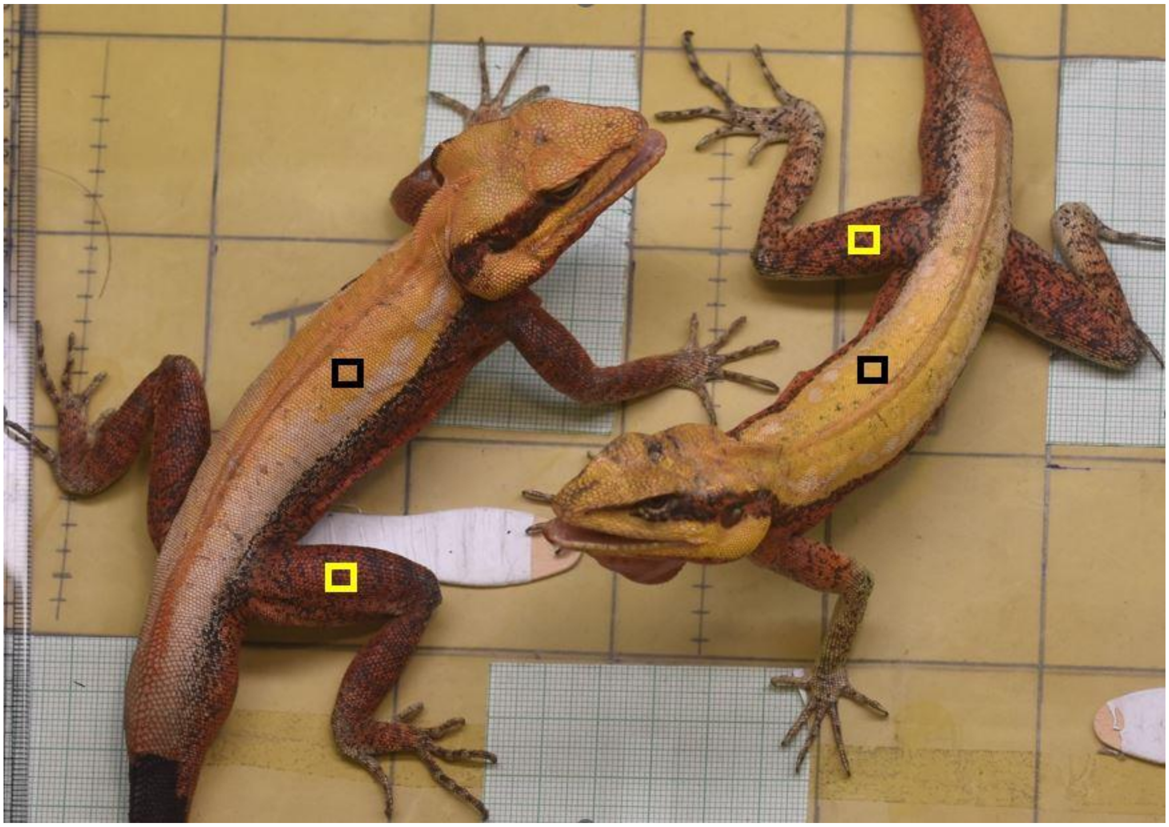

**Figure S1.** Staged male-male interaction in *P. dorsalis* where the black square represents the region of dorsal colour and yellow square represents the region of lateral colour measurement from both the interacting lizards from digital images.

a)

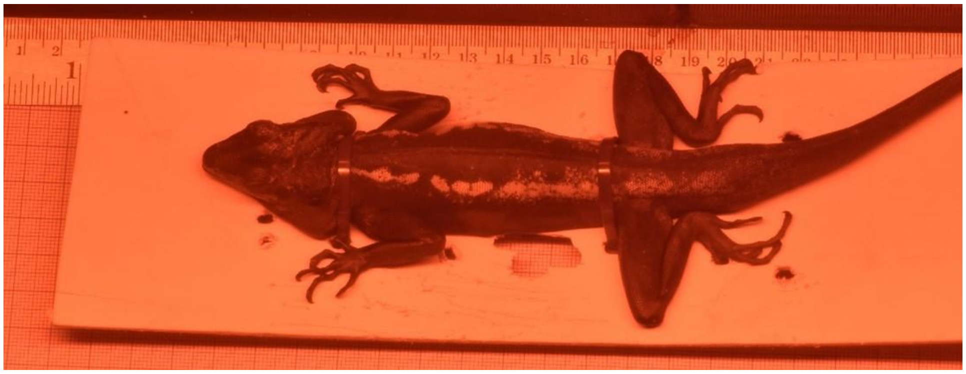

b)

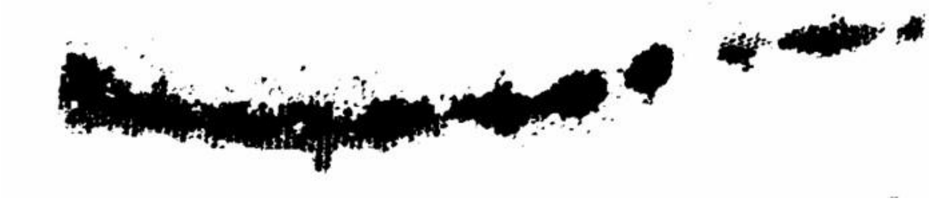

**Figure S2.** (a) UV photograph of a restricted *P. dorsalis* male taken after 10 minutes of stress handling. (b) UV patch extracted from the UV images using the threshold tool in

a)

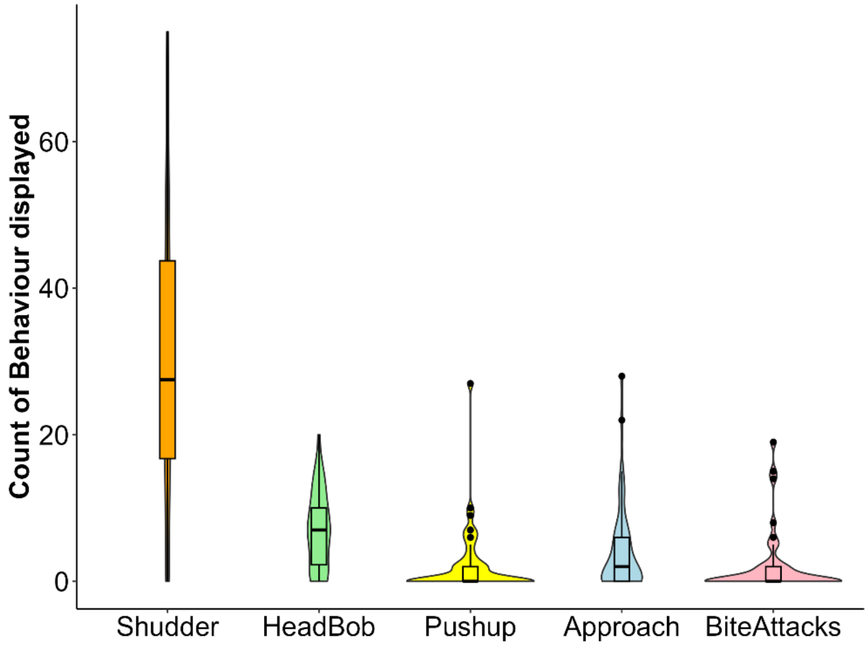


-

b)

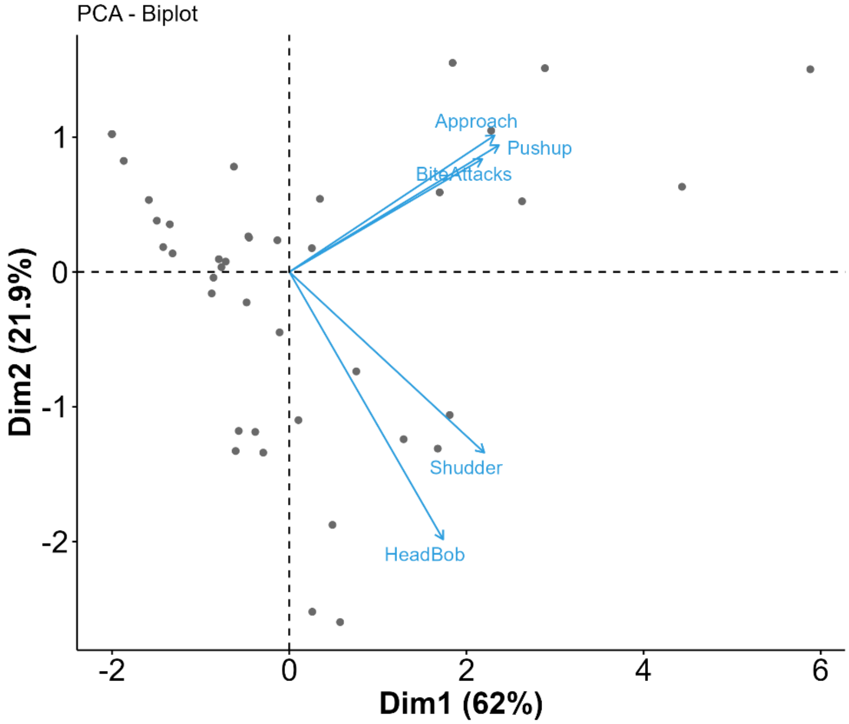


**Figure. S3.** (a) Behaviours displayed by males of *P. dorsalis* males during staged male-male interactions. (b) PCA of the display behaviours with PC1 and PC2 explaining 84% of the data.

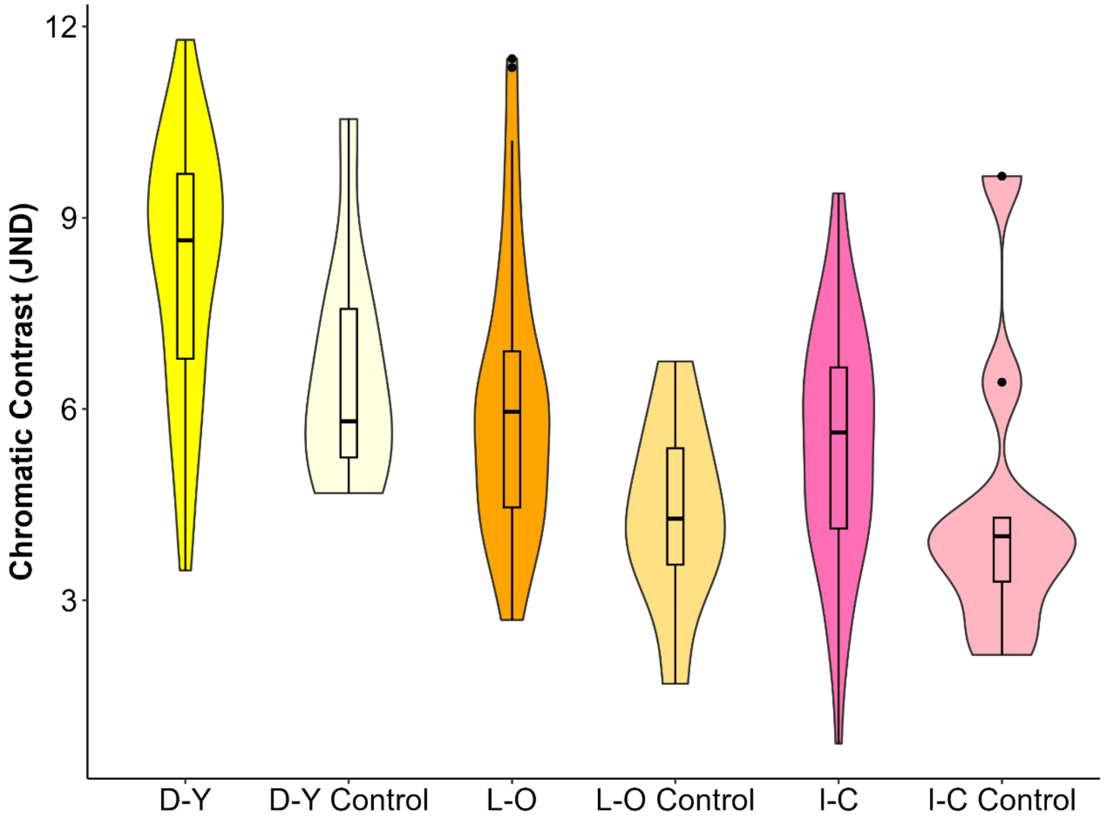


**Figure. S4.** Chromatic contrast of the dorsal yellow (D-Y) and lateral orange (L-O) body regions along with the internal contrast (I-C) between these regions, for males of P. dorsalis that engaged in social interaction and those that did not engage in social interactions (control)

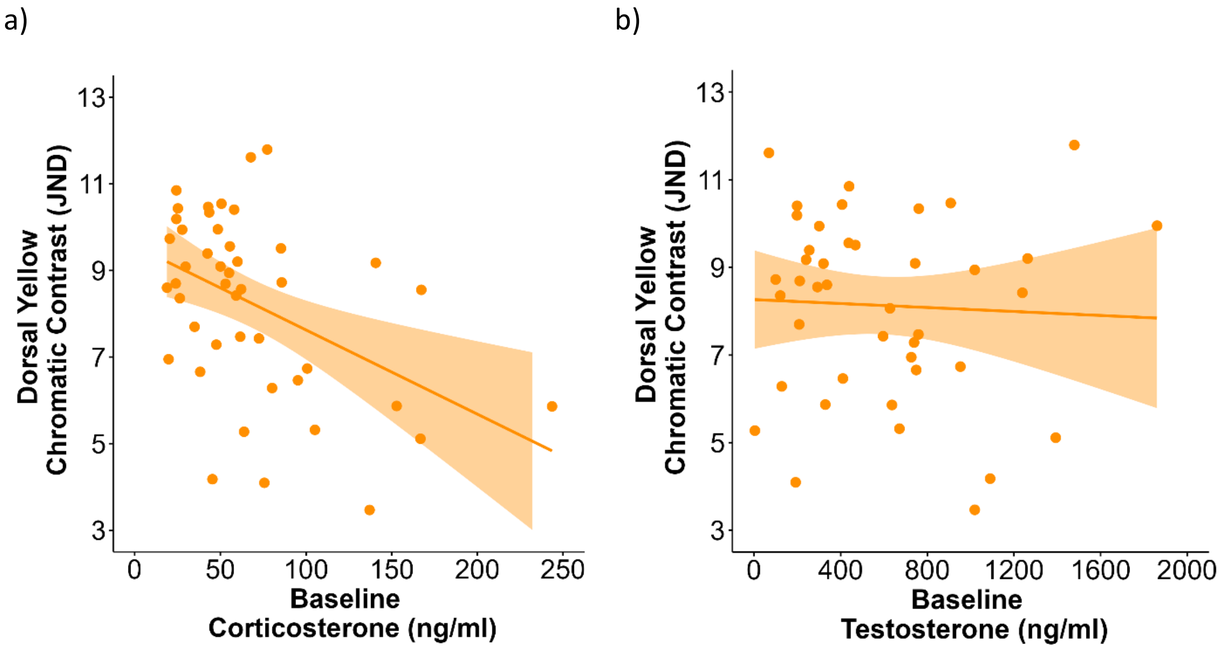


**Figure S5**. Chromatic contrast of the dorsal body region was (a) negatively correlated with baseline corticosterone levels (r = -0.41, p = 0.005) but (b) uncorrelated with baseline testosterone levels (r = -0.05, p = 0.74).

a)

b)

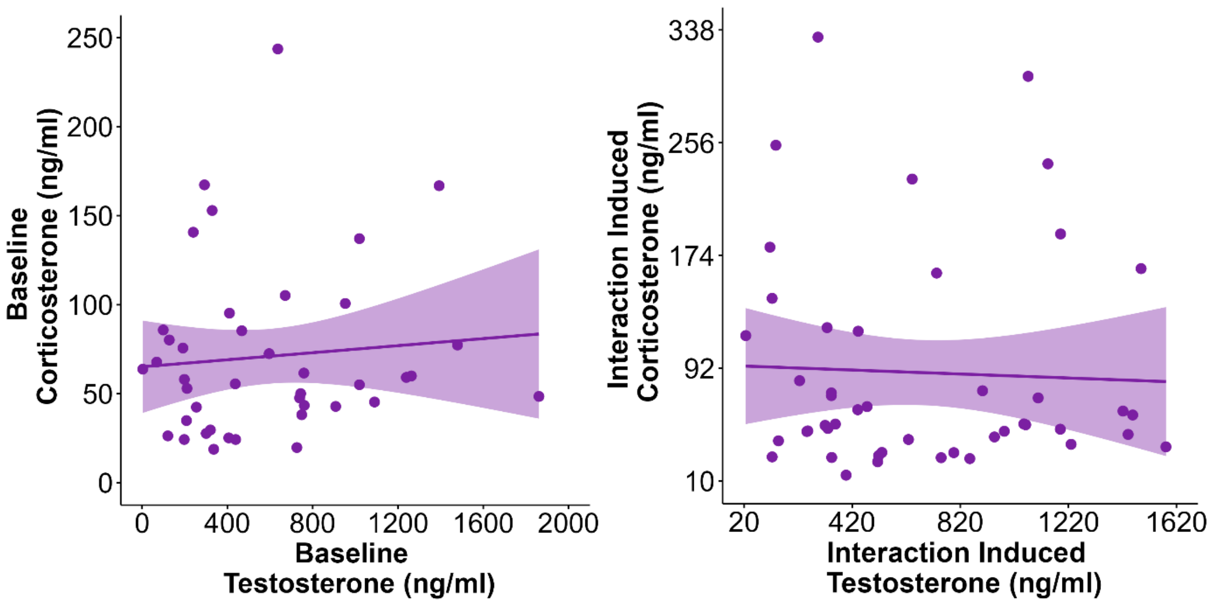


**Figure S6.** (a) Baseline (r = 0.07, p = 0.65) and (b) interaction induced levels (r = -0.09, p = 0.57) of testosterone and corticosterone levels were not correlated with each other.

a)

b)

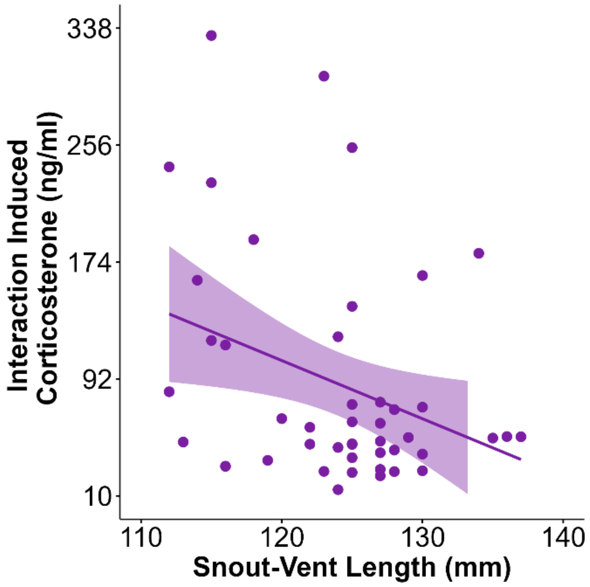


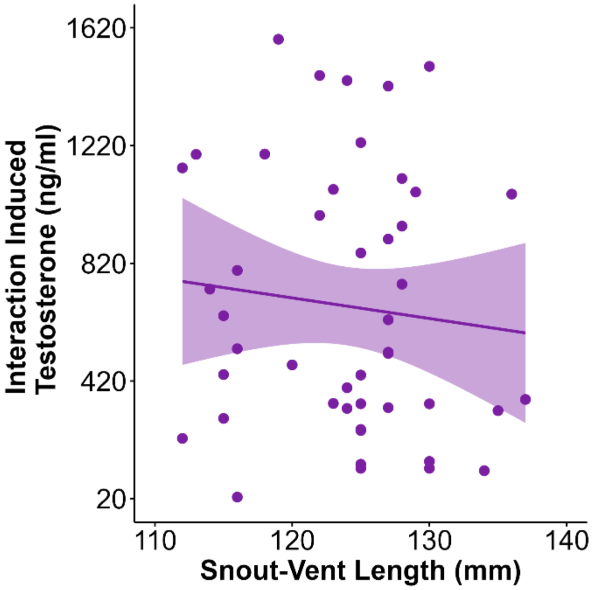


**Figure S7.** Body size (SVL) was not correlated with either (a) interaction-induced corticosterone ($r = -0.24$, $p = 0.1$) or with (b) interaction-induced testosterone ($r = -0.13$, $p = 0.38$).
